# Fluctuation-Driven Synergy, Redundancy, Signal to Noise Ratio and Error Correction in Protein Allostery

**DOI:** 10.1101/2024.12.26.630391

**Authors:** Burak Erman

## Abstract

This study explores the relationship between residue fluctuations and molecular communication in proteins, emphasizing the role of these dynamics in allosteric regulation. We employ computational tools including the Gaussian Network Model, mutual information, and interaction information, to analyze how stochastic interactions among residues contribute to functional interactions while also introducing noise. Our approach is based on the postulate that residues experience continuous stochastic bombardment from impulses generated by their neighbors, forming a complex network characterized by small-world scaling topology. By mapping these interactions through the Kirchhoff matrix framework, we demonstrate how conserved correlations enhance signaling pathways and provide stability against noise-like fluctuations. Notably, we highlight the importance of selecting relevant eigenvalues to optimize the signal-to-noise ratio in our analyses, a topic that has yet to be thoroughly investigated in the context of residue fluctuations. This work underscores the significance of viewing proteins as adaptive information processing systems, and emphasizes the fundamental mechanisms of biological information processing. The basic idea of this paper is the following: Given two interacting residues on an allosteric path, what are the contributions of the remaining residues on this interaction. This naturally leads to the concept of synergy, redundancy and noise in proteins, which we analyze in detail for three proteins CheY, Tyrosine Phosphatase and β-Lactoglobulin.

## 1. Introduction

Evolution has endowed proteins with a remarkable capacity for efficient and robust nanoscale communication and information transmission. This capability is rooted in several fundamental features, including large-scale spatially correlated fluctuations of residues that facilitate dynamic interactions within the protein structure. A network of interactions, characterized by small-world scaling topology(Atilgan, et al., 2004) with spectral characteristics in the Terahertz region(Elayan, et al., 2022), conveniently probed through molecular dynamics simulations, establishing effective pathways for information transfer. Additionally, the synergistic effects of specific residues enhance signaling along these pathways, while redundancy in the fluctuations of certain residues provides stability and acts as a corrective mechanism to ensure reliable signal propagation. Together, these factors contribute to the overall resilience and efficiency of intramolecular signal transmission in proteins, highlighting their role and functional significance in allosteric processes (Blacklock and Verkhivker, 2014; Hamborg, et al., 2021; Yang, et al., 2023). The spatial and temporal replication of the same protein with high precision across all species clearly demonstrates that the aforementioned fundamental features are universal determinants of protein function (Tang and Kaneko, 2021).

The network topology of a protein is mathematically represented by the Kirchhoff matrix, K, a computational construct that transforms our understanding of molecular communication. Defined as Γ = D - A, where D is the degree matrix and A is the adjacency matrix, this framework captures protein residue connectivity. Computational validation studies have demonstrated that these matrices can map intricate molecular communication networks that traditional methods overlook (Bahar, et al., 1997). The inverse of the Kirchhoff matrix, Γ⁻¹, generates the covariance matrix Σ, a dynamic representation of molecular motion that gives residue fluctuation correlations (Haliloglu, et al., 1997). Biological validation studies have shown that these matrices can predict functional mutations with up to 70% accuracy, bridging theoretical models with experimental observations (Mishra, 2022; Zheng, 2024). By employing a linear spring model, we can describe molecular movements through a multivariate Gaussian probability density function. This probabilistic approach forms the basis of information theory of proteins that we adopt here (Hacisuleyman and Erman, 2022).

Entropy quantifies the dynamic information transfer within proteins, playing a crucial role in allosteric regulation. It reflects how conformational changes at one site can influence distant regions, facilitating communication essential for protein function. This entropic interplay enables proteins to adaptively respond to stimuli, allowing for precise control over biological processes (Cooper and Dryden, 1984; Hacisuleyman and Erman, 2017).

Mutual information provides a nuanced understanding of residue interactions that predicts functional relationships between residues with remarkable computational accuracy. Computational studies have shown that residues with high mutual information are often involved in critical cellular processes, such as signal transduction and enzymatic catalysis. In G-protein coupled receptors, for example, mutual information analysis has uncovered communication pathways explaining how distant mutations can dramatically alter protein function (Demirtaş, et al., 2024).

Conditional mutual information represents an advanced approach to understanding molecular communication mechanisms. This metric has been particularly powerful in explaining allosteric phenomena, such as hemoglobin’s cooperative binding—where oxygen binding to one subunit induces conformational changes in distant protein regions, demonstrating the complex, interconnected nature of protein dynamics (Daura, 2019; LeVine and Weinstein, 2014).

Interaction information emerges as a sophisticated tool for understanding molecular relationships beyond simple pairwise interactions. Positive interaction information suggests redundancy, potentially representing backup pathways and molecular error-correction mechanisms, while negative interaction information reveals synergistic communication pathways (Hacisuleyman and Erman, 2024).

Interactions between residues are inherently stochastic, either enhancing each other’s functional integrity or contributing to noise. Each residue experiences continuous stochastic bombardment from impulses generated by all other residues, whether they are nearby or distant, depending on the connectivity of the interaction network topology. A subset of these impulses consists of evolutionarily conserved and essential correlations for the protein’s function and stability, while the remaining impulses are random and noise-like (Erkip and Erman, 2004; Ormos, 2008; Tang, et al., 2020; Zheng and Tekpinar, 2009). In either case, the information-theoretic tools we employ in this paper shift our perspective and treatment of proteins from stochastic molecular structures to adaptive information processing systems, especially regarding their spectral properties. Notably, while the signal-to-noise ratio in these analyses depends strongly on the number of eigenvalues considered, highlighting the importance of selecting relevant modes for accurate predictions, the specific problem of signal-to-noise ratio related to residue fluctuations has not yet been thoroughly investigated. Recent breakthrough studies have utilized these methods to design novel proteins with enhanced stability and to develop targeted therapeutic interventions aimed at understanding the molecular basis of genetic diseases (Dantas, et al., 2003; Leaver-Fay, et al., 2011). By mapping intricate information networks, the present study paves the way for predicting mutation-induced communication disruptions, elucidating protein misfolding mechanisms, and advancing sophisticated protein engineering strategies. The implications extend far beyond basic structural biology, offering significant insights into cellular signaling, drug design, and the fundamental mechanisms of biological information processing.

In this paper, we utilize three well-studied cases of allosteric activity, CheY, Tyrosine Phosphatase and β-Lactoglobulin. We use the Gaussian d Network Model, GNM, to demonstrate the role of residue fluctuations in information transfer and to explain the strategies employed to achieve synergy and redundancy in allosteric communication.

## 2. Materials and Methods

### 2.1 Allostery in proteins and metrics of information theory

The problem that we are interested in is the following: We assume that two residues, say A and B, in a protein interact, and this interaction is essential for allosteric activity. If the two residues are within the coordination shell of each other, they will interact through typical Lennard-Jones forces, including hydrogen bonds, electrostatic interactions, and steric effects. They may be spatially distant. In this case, the graph structure of the protein significantly influences their interaction (Clementel, et al., 2022; del Sol, et al., 2006).

If a third residue C enhances the interaction between A and B, its effect is considered synergistic, forming what we term the allosteric channel. Conversely, C may provide overlapping information about the interaction between A and B that is already conveyed by either A or B. In information theory, this overlapping information is referred to as “redundant.” However, in the context of proteins, C can be viewed as a backup residue that supports the A-B interaction, particularly if that interaction is weakened due to mutations or dynamic changes within the protein. Therefore, we will use the term “redundant information” to mean “overlapping information.” The residues that provide this information will be called “redundant residues,” but not in a negative sense.

Entropy of a residue A is represented by the Shannon equation (Cover, 1999)

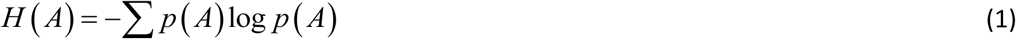

where, A represents the set of fluctuations of residue A, p(A) is the probability and H(A) quantifies the degree of uncertainty associated with those fluctuations. For correspondence between the thermodynamic entropy and the Shannon entropy, please see Chapter 17 of the book by Callen (Callen, 1985).

Mutual information, *I* (*A*; *B*) between A and B is defined as

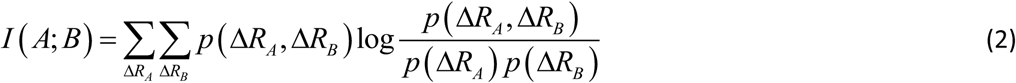

where, *p* (Δ*R_A_*, Δ*R_B_*) is the joint probability of fluctuations of residue A and B. The summation is carried out over all fluctuations of A and B. For brevity, the notation will be simplified by replacing *p* (Δ*R_A_*, Δ*R_B_*) with *p* (*A*, *B*) and *p* (Δ*R_A_*) with *p* (*A*), etc., in the following. *I* (*A*; *B*) quantifies the reduction in uncertainty in the fluctuations of B that is achieved by knowing the fluctuations of A. If A and B are independent, meaning that knowing A gives no information about B, the mutual information is zero. However, if there is some relationship—whether linear or non-linear—mutual information will be positive.

Joint entropy *H* (*A*, *B*) is defined as

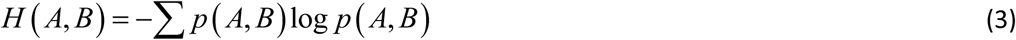

Conditional entropy *H* (*B* | *A*) is defined as

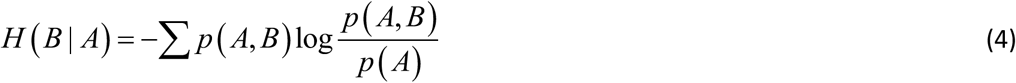

*H* (*B* | *A*) measures the amount of uncertainty remaining in the fluctuations of residue B when the fluctuations of residue A are known. It quantifies how much additional information is needed to describe B given that we already have information about A. When we observe the fluctuations of residue A, we may gain insights into the behavior of residue B. If knowing the state of A significantly reduces our uncertainty about B, then H(B∣A) will be low. This indicates that the fluctuations of A and B are correlated; when A fluctuates, B tends to fluctuate in a predictable manner.

Mutual information is related to entropy by the equation

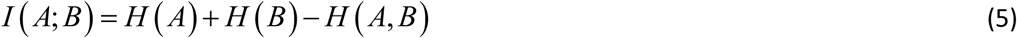

Conditional mutual information in terms of entropy is

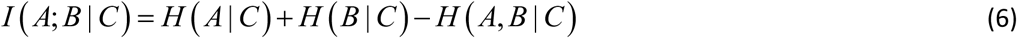

Conditional mutual information measures the amount of information that two random variables, A and B, share in the presence of a third variable, C. And finally, the interaction information

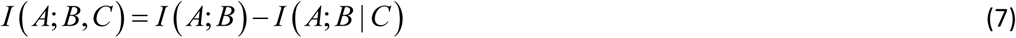

which represents a measure of how the information shared between variables A and B changes under the effect of another variable C. If the presence of C decreases the interaction between A and B, then *I* (*A*; *B*) > *I* (*A*; *B* | *C*) and *I* (*A*; *B*, *C*) is positive. The amount of information provided by C to the interaction between A and B in this case has an overlapping component. The lowering of *I* (*A*; *B* | *C*) due to the presence of C may be due to two different effects that may be operating in the system. First, the presence of C may create mechanical noise, which may affect the direct correlation between A and B. Calculations show that the noise created by neighboring residues C along the primary chain increase the interaction information between A and B. The second effect occurs when C acts similarly to A or B in their interaction. In this case, C provides overlapping or duplicate information. This causes the conditional mutual information *I* (*A*; *B* | *C*) to decrease because C does not add unique information but rather repeats what A and B are already doing. This second case may indeed be useful for the protein to carry its allosteric function. It will be needed when the protein experiences mechanical stress, when one allosteric pathway is compromised, helps to ease-down negative effects of mutation, provides robustness against genetic variations, creates alternative information transmission routes, compensates for local structural fluctuations, maintains interaction integrity under perturbations, and prevents catastrophic information loss. The key point is that redundancy occurs when an additional element (C in this case) provides information that is already largely captured by the existing interaction between A and B. It is interesting to note that both redundancy and noise can lead to similar mathematical outcomes (positive interaction information), but they represent fundamentally different processes: Redundancy strengthens or confirms existing relationships. Noise, on the other hand, introduces uncertainty and irrelevant information.

If the presence of C increases the interaction between A and B, which we call synergy, then *I* (*A*; *B*) < *I* (*A*; *B* | *C*) and the interaction information is *I* (*A*; *B*, *C*) is negative. By identifying synergetic residues, it becomes possible to map potential allosteric paths. Such synergetic residues are essential in understanding long-range communication mechanisms.

The interaction between A and B is subject to impulses from all other residues of the protein. The information received by A-B is *I* (*A*; *B*,*C*) where C includes all residues other than A and B. The standard deviation, SD, of the information that comes to A-B is representative of the information transferred by C. Given this description of the signal (I(A;B,C)), the noise naturally comes from the standard deviation of the conditional mutual information, SD(I(A;B|C)) showing the variability or uncertainty in the information transfer between A and B when accounting for all other residues. This standard deviation quantifies the fluctuations and inconsistencies in how the other residues modulate the information exchange between A and B. Higher standard deviation indicates more unpredictable or scattered information transfer, suggesting less stable or more complex interactions between A and B in the context of the entire protein. Thus, the signal to noise ratio, *S* / *N* becomes

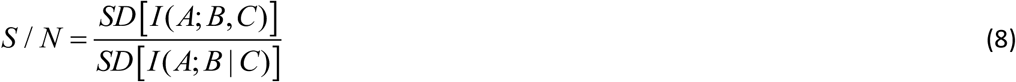

The signal-to-noise ratio (S/N) strongly depends on the spectral properties of the protein, particularly with larger eigenvalues of the Kirchhoff matrix being more representative of noise, while noise-like eigenvalues contribute more significantly to the denominator of Eq. 8.

Intramolecular noise provides insights into the robustness and adaptability of allosteric pathways, highlighting how internal redundancy or synergy can maintain effective communication despite perturbations. This perspective aligns with the principles of information theory, where variability in pathways can serve as a basis for error correction and signal reliability.

Finally, we consider the error correction capability of C. C provides overlapping information about the interaction between A and B, where transmission errors may occur. Here, we use the term “transmission” in its broadest sense. For example, a sporadic impulse that tends to break a necessary hydrogen bond between A and B may be regarded as a transmission error that can be masked by the information sent by C. In this context, C acts as a molecular error correction code. Alternatively, if the A-B interaction is compromised or weakened, the overlapping information provided by residue C may reduce the uncertainty introduced. This suggests a natural error correction mechanism in protein communication pathways that compensates for imperfections in primary signal transmission and restores the integrity of the signals.

### 2-2. The Gaussian Network Model and information transfer

The Gaussian Network Model (GNM) is a coarse-grained representation of proteins, where each Cα atom is modeled as a node in a graph, and edges connect nodes that are either covalently bonded or spatially proximate. The graph’s connectivity is described by its Laplacian or Kirchoff matrix, Γ, which assigns unit entries to connected nodes and diagonal elements equal to the negative sum of each row. This Laplacian encodes the protein’s local structural features, i.e., the residue interaction matrix, while its inverse is proportional to the fluctuation correlation matrix

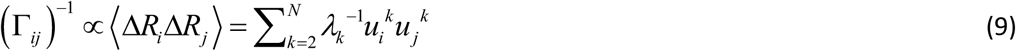

where, angular brackets denote average, i and j refer to residue indices, *λ_k_* is the k’th eigenvalue of Γ, and*u_i_ ^k^* is the k’th eigenvector of Γ for the i’th residue. The model approximates fluctuations as isotropic, and despite this simplicity, it remarkably aligns with experimental observations of protein dynamics (Bahar, et al., 1997).

The inversion of the Laplacian matrix shows a key strength of GNM: it transforms local structural information into a global interaction map. In the original Laplacian, residue pairs that are physically distant are represented with zero entries due to the lack of direct interactions. However, the inverse Laplacian introduces nonzero entries for these pairs, capturing the indirect interactions mediated through the network, which automatically contributes to the mutual information between distant residue pairs. These nonzero values reflect dynamic couplings between residues, offering insight into long-range relationships within the protein. This capability to account for allosteric interactions in a single computational step distinguishes GNM from other methods.

In addition to providing structural insights, GNM captures protein dynamics by analyzing the eigenmodes of motion derived from the Laplacian (Haliloglu, et al., 1997). These eigenmodes explain collective movements and pathways that underlie functional processes such as allosteric signaling. The Laplacian has a zero eigenvalue, which corresponds to rigid body motion. Smaller eigenvalues characterize slow and large scale motions and larger eigenvalues correspond to fast motions that are more noise-like. By integrating both structural and dynamic information, GNM offers a computationally efficient framework for studying information transfer and long-range communication within proteins, making it the simplest yet powerful approach for such analyses.

Entropy for n variables for a multivariate Gaussian distribution of fluctuations reads as

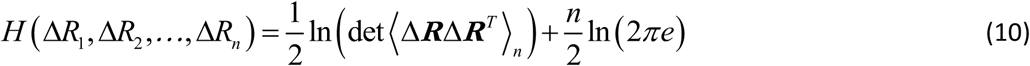

Expressions for mutual information, conditional mutual information and interaction information are obtained from the entropy using equations 3-7.

Substituting from Eq. 10 leads to the following

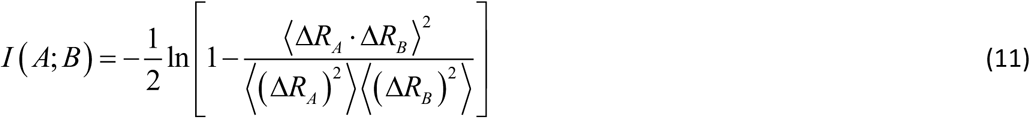

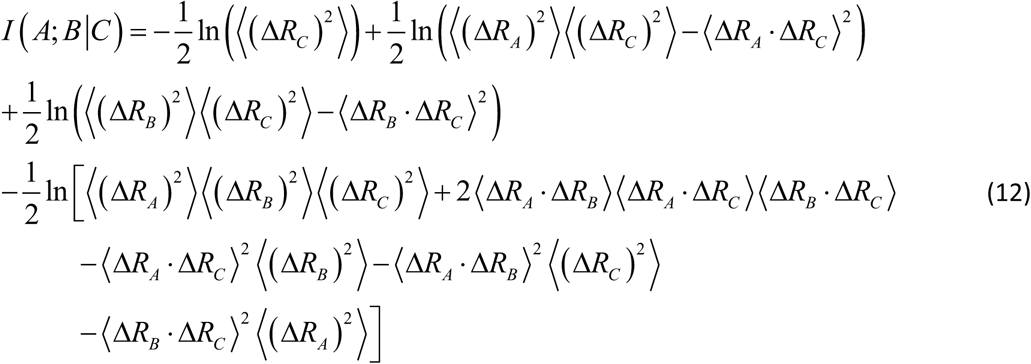

All entries in Eqs. 11 and 12 follow from the inverse of the Laplacian and their use in the interaction information expression, Eq. 9. Scanning over the protein by keeping A and B fixed and changing C leads to the interaction information profile for the protein in which positive peaks show the backup residues that provide overlapping information and negative peaks show the synergistic residues for the interaction between residues A and B.

When C equals A or B, then the right hand side of Eq 7 is undefined, but since *I* (*A*; *B* | *B*) = 0 we have

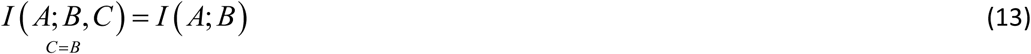

## 3. Results

### 3-1. CheY bacterial chemotaxis

CheY is a signaling protein in the bacterial chemotaxis signaling pathway. When phosphorylated at D57, the protein travels to a large flagellar motor complex and binds to FliM, which is part of the motor complex. When CheY binds to FliM, it causes the switch complex to change its rotation direction. The primary effect is the increased interaction of D57 with T87, enabling CheY to bind to FliM. The interaction is in the form of a hydrogen bond which forms when D57 is phosphorylated. It helps maintain the allosteric activity effectively. Thus, the allosteric mechanism of CheY occurs through changes in its dynamical properties upon the interaction of D57 with T87 rather than a simple, mechanistic structural rearrangement. The problem we investigate here is the following: Given the interaction of D57 and T87, how do the remaining residues of the protein affect this interaction. We are particularly interested in the remaining residues of the protein that enhance this interaction or serve as a backup node for the maintenance of this interaction. We also investigate the level of information and noise coming to this interaction.

#### 3-1-1. Determining the synergistic and overlapping residues controlling the interaction between D57 and T87

The interaction information profile for CheY, calculated according to Eqs. 7, 11 and 12 for *I* (57;87, *C*) where C represents each residue scanned across the entire protein is presented in Figure 1. The negative synergistic and positive redundant peaks in the figure are supported by experimental observations (Cho, et al., 2000; McDonald, et al., 2012).

**Figure 1.**
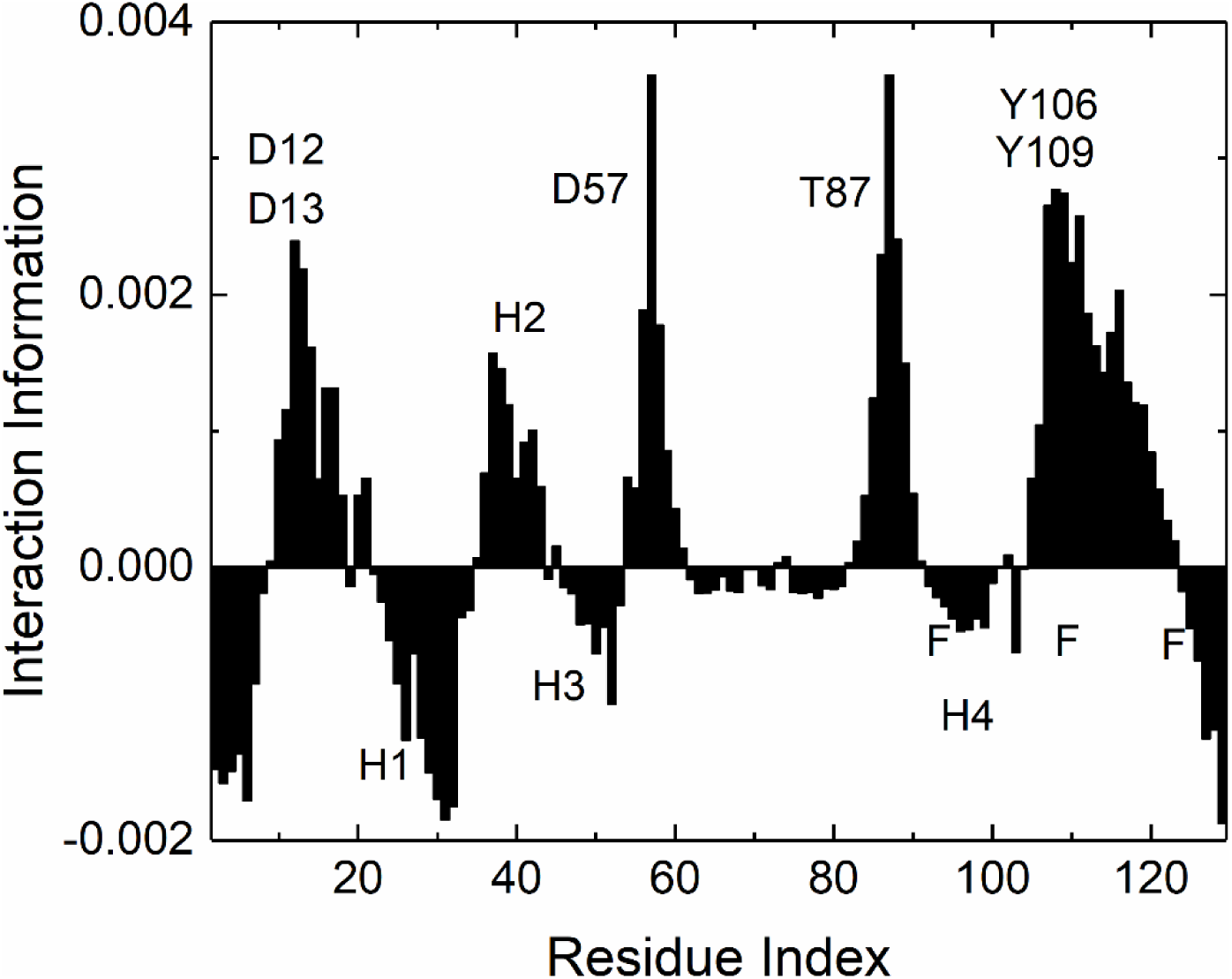
Interaction information *I* (57;87, *C*) where C is scanned over all residues of the protein shown along the abscissa. The Protein Data Bank structure 1F4V is used in the calculations with a cutoff distance of 7.3 Å in obtaining the Γ matrix. The proportionality constant for Eq. 9 is taken as unity. All modes are used in the Negative peaks show the synergistic contribution of the corresponding residue to the interaction between D57 and T87. Positive peaks correspond to residues that provide overlapping information to this interaction. D12, D13, D57, T87 and L106 and K109 are conserved residues(Lowry, et al., 1994). Binding sites of Flim are indicated by F. Helices are indicated by H. Interaction information values are obtained keeping all modes of the Kirchoff matrix.

**Figure 2.**
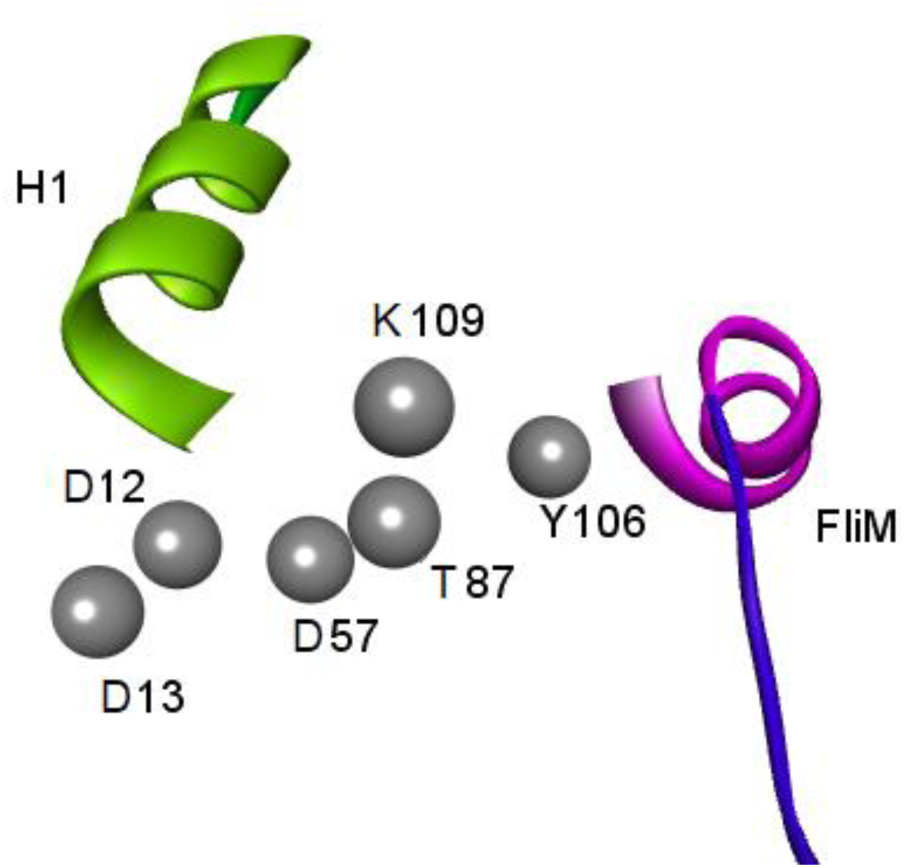
The allosteric path obtained from peaks of Figure 1. Alpha carbons of the six conserved residues are shown with gray spheres. Allosteric activity starts at D57 when it is phosphorylated by the protein CheA. This is followed by the rearrangements of the conformations of the residues shown such that FliM becomes able to bind to CheY as a result of which the flagellar motor attached to FliM is activated. Helix1 and the conserved residues shown are derived from the peaks of the interaction information profile *I* (57;87, *C*), which is calculated for each residue of CheY.

The conserved D12 and D13 residues, though located on the opposite side of the protein from D57 and the FliM binding site, i. e., residues A90, K91, K92, G105, Y106, and K109, contribute redundant information to the D57-T87 interaction and act as backup nodes. This means that they can compensate for the primary interaction pathway, can help maintain the functional relationship between D57 and T87 even if the original interaction is compromised and can provide an alternative route for information, or signal transmission. Both D12 and D13 coordinate the binding of the Mg(II) ion, which is necessary for the activation of CheY. This ion is essential for stabilizing the structure of CheY and facilitating its interaction with other proteins during allosteric activity (Bourret, et al., 1993).

The first 10 residues of the protein act synergistically on the interaction of D57 and T87 and are critical for establishing the overall structure of CheY. They help define the protein’s conformation, which is essential for its interaction with other proteins and for transmitting signals upon phosphorylation. Phosphorylation of D57 is a conditional requirement for changes in the conformations of the first ten residues, modulating CheY’s interaction with the flagellar motor. Mutations in these residues are shown to lead to altered conformations that affect CheY’s ability to activate flagellar rotation (Bourret, et al., 1993; Quax, et al., 2018).

The first helix, H1, is the strongest synergistic region strengthening the interaction between D57 and T87. The formation of hydrogen bonds between residues in helix 1 and the neighboring beta strand contributes to the overall stability of CheY which is the source of communication between H1 and D57 (Wilcock, et al., 1998). According to Figure 1, the second helix, H2, acts redundantly and the third helix, H3, acts synergistically on the interaction between D57 and T87. Previous work (Haliloglu, et al., 2022; Wang, et al., 2020) identified the importance of the helices H1-H3 in allosteric communication. The present model is able to differentiate the mechanism of action between these helices as to whether they are synergistic or redundant. Synergy amplifies effects and leads to emergent properties in signaling pathways, while redundancy tends to maintain baseline functionality despite perturbations.

Mutual information between D57 and T87 is represented by the two vertical lines at positions 57 and 87 in Figure 1. The positive peaks in the neighborhood of D57 and T87 show that these residues provide overlapping information to the 57-87 interaction.

The nature of the information transmitted from D57 to FliM can be visualized in three distinct forms:

a. Biochemical State Information: Phosphorylation at D57 alters the charge and chemical properties of this residue. This modification serves as a biochemical signal that switches CheY from an inactive to an active state.
b. Conformational Information: The phosphorylation event induces a conformational shift in the structure of CheY, particularly in residues surrounding the phosphorylation site, including D12 and D13. These residues play a critical role in stabilizing the phosphorylated state and facilitating the propagation of the activation signal to CheY’s FliM-binding interface, as illustrated by F in Figure 1. This process effectively “transmits” information about CheY’s activation state.
c. Functional Information: The final “output” of this information transfer is CheY’s ability to bind FliM. The phosphorylation and subsequent conformational changes, supported by D12 and D13, create a high-affinity binding interface for FliM, determining whether CheY can influence the rotational state of the flagellar motor (clockwise or counterclockwise).

Explaining the allosteric activity of CheY in terms of information theory adds a new dimension to our previous allostery analysis of this system (Hacisuleyman and Erman, 2022).

#### 3-1-2. Effect of noise on information transmission in CheY and the signal to noise ratio, SNR

The effect of a third residue on the interaction between a given pair of residues depends strongly on the number of modes of Eq. 9 used in the analysis. As the number of modes increases, the magnitudes of the interaction information profiles tend to decrease significantly because additional modes contribute predominantly to noise rather than meaningful interactions. This dilution of signal obscures the significance of the main interactions in the allosteric process. However, the shape of the profile remains approximately independent of the number of modes used, showing that residues contributing to synergy and redundancy are identified irrespective of the number of modes. In Figure 3, profiles are shown for four different cases where 10, 20, 50 and all modes are used in calculations. One sees that while the general shape remains approximately the same for the four different cases, the amplitudes drop down significantly as the number of modes becomes larger.

**Figure 3.**
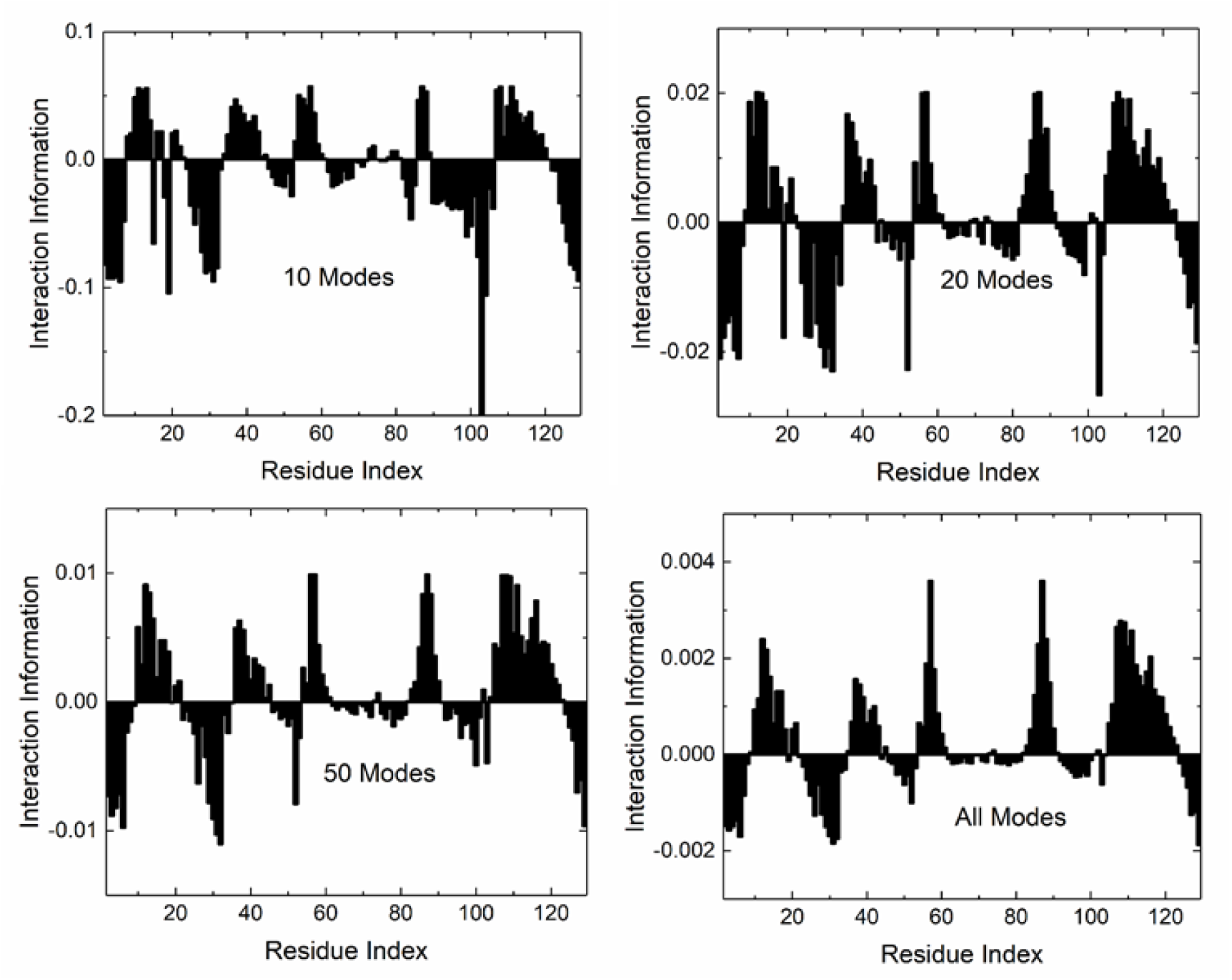
Interaction information profiles obtained by using different number of modes in Eq.9. The number of modes used is shown in each panel.

The question of ‘how many modes should be used for a representative case’ can be answered by calculating the signal-to-noise ratio (SNR) as a function of the number of modes, as shown in the following figure.

**Figure 4** illustrates that the use of the lowest eight modes results in signal-to-noise ratio (SNR) values greater than 0.5, which can be accepted as contributing to the allosteric signal affecting the interactions between residues D57 and T87. It is reasonable to conclude that higher modes correspond to such small amplitudes that they collectively contribute to noise in the context of information propagation. An order of magnitude analysis of the mean-square fluctuations of CheY residues, along with their decomposition into modes and comparison with experimental B-factors, indicates that the first ten modes are associated with displacements greater than approximately 0.1 angstroms, while higher modes correspond to displacements below this threshold.

**Figure 4.**
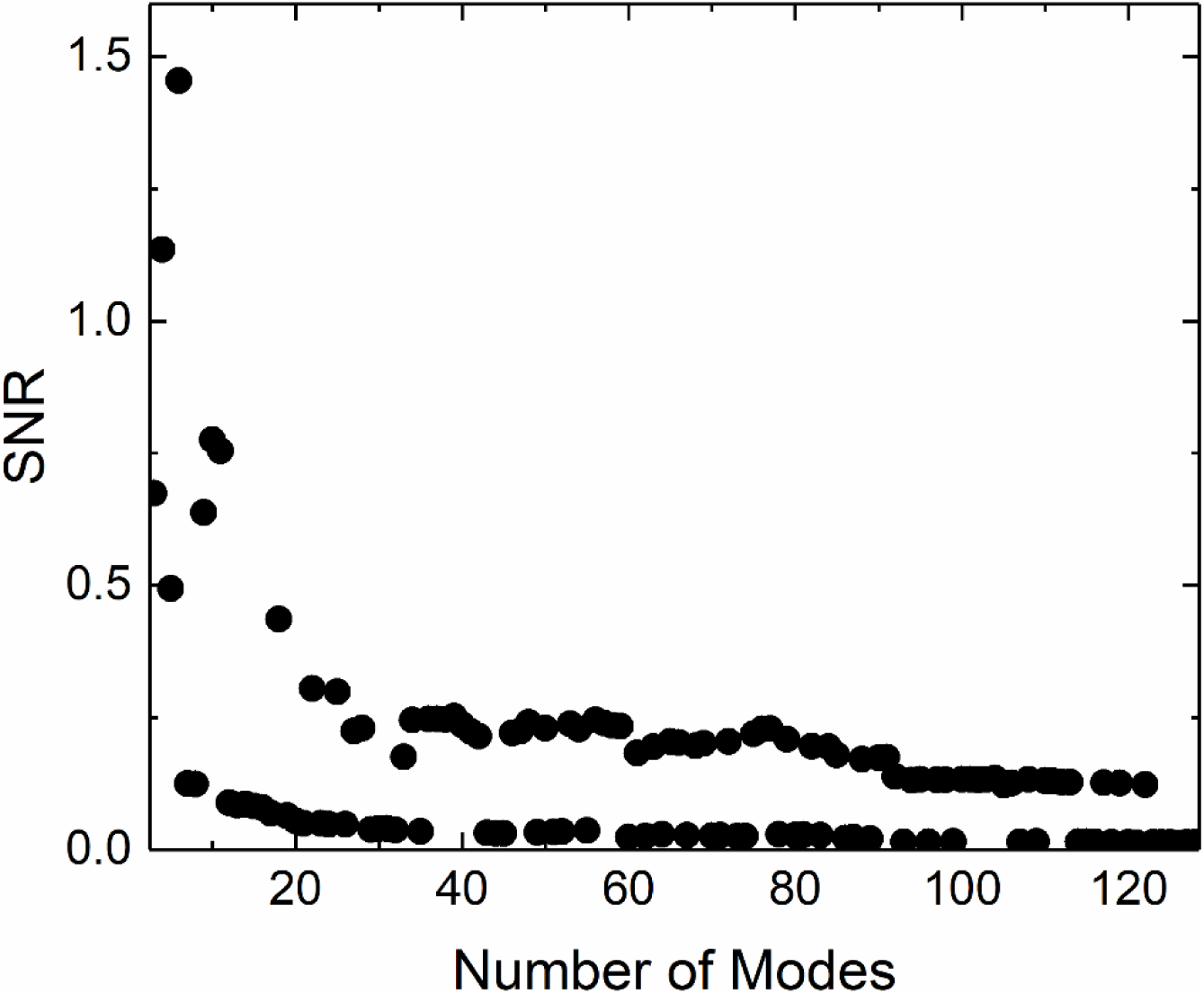
SNR graph for CheY. The points are obtained using Eqs. 11 and 12 for the number of modes shown along the abscissa. See legend to Figure 1 for details of calculation.

It is interesting to note that the plot of SNR versus the number of modes shows significant fluctuations.

While using *n* modes results in a high value for information, the SNR decreases drastically when *n*+1 modes are used. Closer inspection of the results shows that these abrupt changes occur when the added mode has an unusually high or low variance in conditional mutual information or interaction information.

#### 3-1-3. Error correction by redundant residues

Finally, in a rather heuristic manner, we discuss a hypothetical but plausible error correction mechanism in CheY, facilitated by redundant residues that support the interaction between residues D57 and T87. These redundant residues provide overlapping information that can stabilize the interaction between D57 and T87, which may be perturbed by structural changes such as mutations or ligand binding. A residue, denoted as C, can help maintain the interaction by compensating for any loss of stability when it fluctuates in a manner that is positively correlated with D57 and/or T87. This means that when D57 and T87 are experiencing unfavorable conditions, residue C should ideally be in a state that supports their interaction. A strong positive correlation between C and these residues can help preserve the necessary interactions or structural integrity despite perturbations caused by mutations or other factors.

In CheY, there are three redundant peaks at residues D12 or D13, E37, and Y109. In Figure 5 we present the sum *v* (*C*, 57) + *v* (*C*,87) of covariance of residue C with both D57 and T87, where the covariances are calculated for different residues C across the entire protein. Notably, we observe that the two redundant peaks, D12/D13 and Y109, exhibit strong positive correlations with both D57 and T87. In contrast, the third redundant peak, E37, which is a smaller redundant peak, does not show a significant positive correlation with D57 and T87.

**Figure 5.**
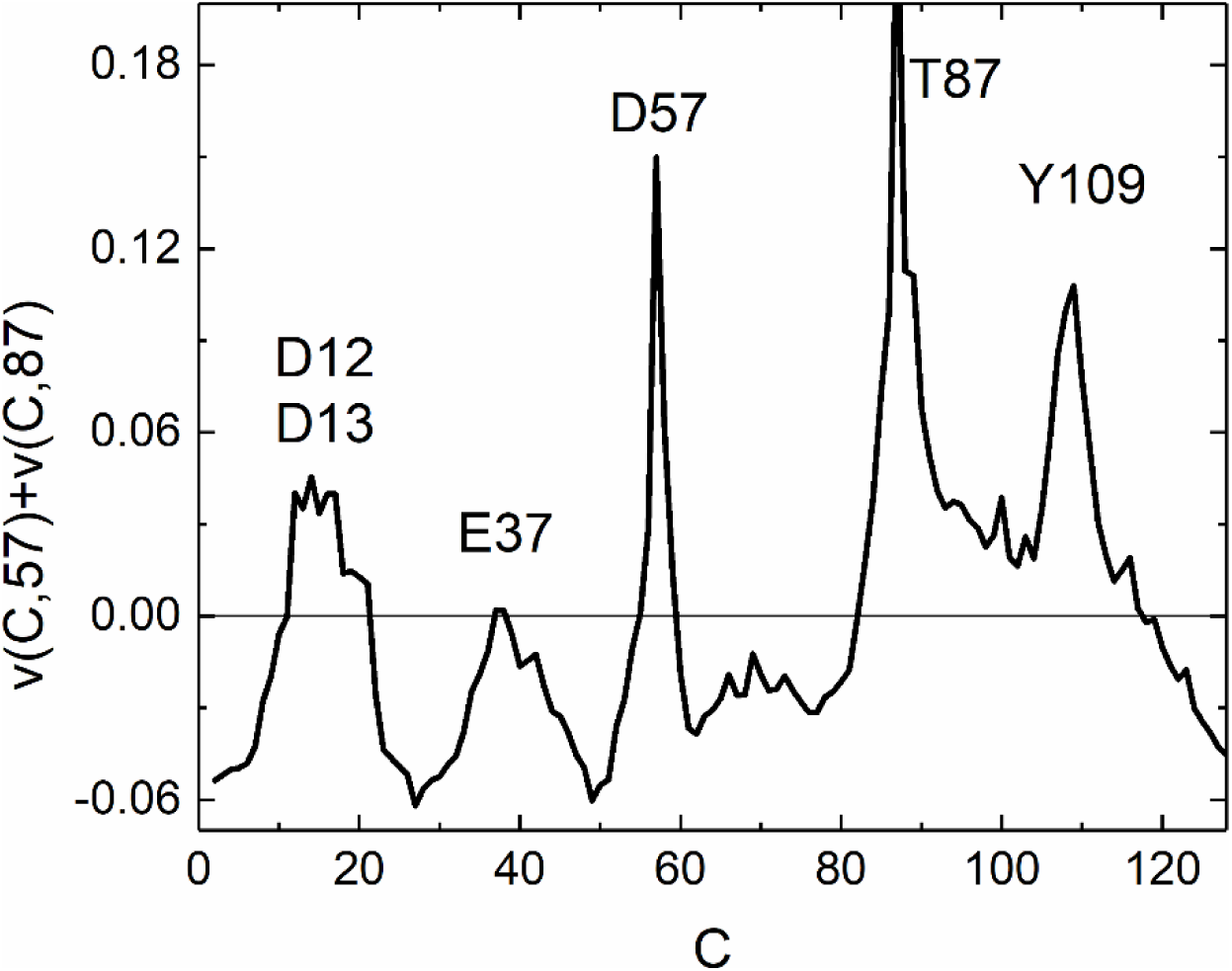
Sum of correlations of C with D57 and T87 for different values of residue C. The correlations are obtained from Eq. 9. See legend for Figure 1 for parameters used.

### 3-2. Tyrosine phosphatase

Tyrosine phosphatase 1B (PTP1B) is a negative regulator of insulin receptor phosphorylation and signaling which is a therapeutic target for type 2 diabetes (Sharma, et al., 2023). Its allosteric activity is described in detail (Hardy and Wells, 2004).

Two residues whose interaction is crucial for allosteric activity of PTPB1 are W291 and L192. In this section, we find residues of PTP1B that effect this interaction. PTP1B has eight helices. W291 is located within helix H7 and is crucial for the allosteric modulation of PTP1B. When an allosteric inhibitor binds to the enzyme around H7, it can displace W291, which disrupts the hydrogen bond network essential for the closure of the WPD loop (residues 179-181) necessary for catalytic activity. This displacement keeps the WPD loop in an “open” conformation, thereby inhibiting the enzyme’s ability to catalyze dephosphorylation reactions (Li, et al., 2014).

Leu192 is situated in helix H3 and plays a significant role in stabilizing interactions with allosteric inhibitors. Its position helps maintain the protein remain in its inactive conformation, preventing the transition of PTP1B to its active form.

Results of calculations using Eqs. 11 and 12 show that PTPB1 consists mostly of backup residues, as presented in Figure 6. Helix 2 is known to connect to the allosteric site, and its conformation is linked to the movement of the WPD loop keeping the WPD loop in an open state upon binding of an allosteric inhibitor. Figure 6 shows that this effect takes place by the synergistic action of H2 on W291 and L192.

**Figure 6.**
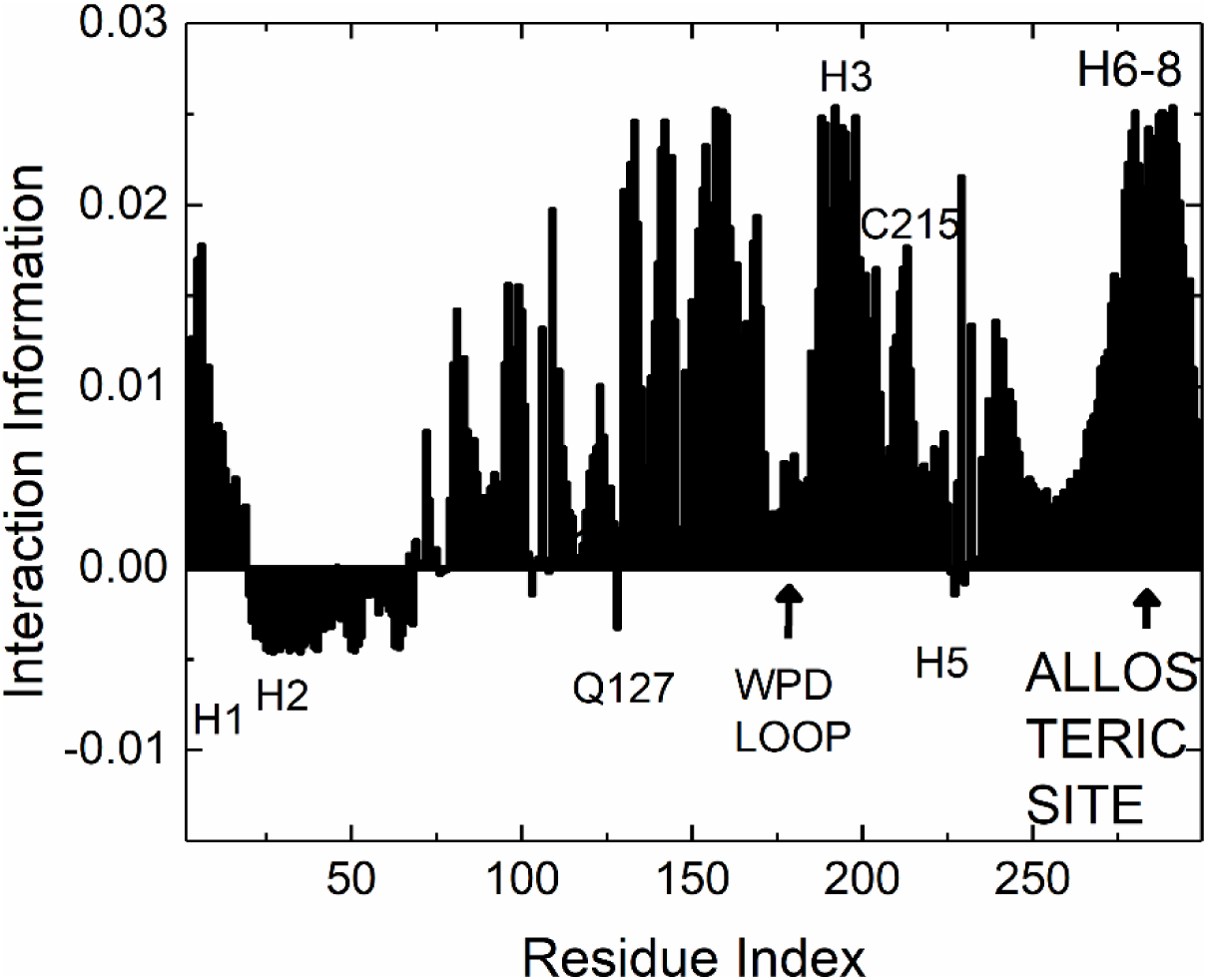
Interaction information profile for PTPB1 for the interaction between W291 and L192. The points are calculated using 60 modes of Eq. 9 (Se explanation in Figure 7). See legend to Figure 1 for details of calculations.

Q127 is in the catalytic site, which also has synergistic effect on the interaction of W291 and L192. In a similar way, the catalytic residue C215 has a synergistic effect on the W291 and L192 interaction.

Signal to noise ratio for PTPB1 is presented in Figure 7 which shows that first 60 modes give SNR values larger than 1.

**Figure 7.**
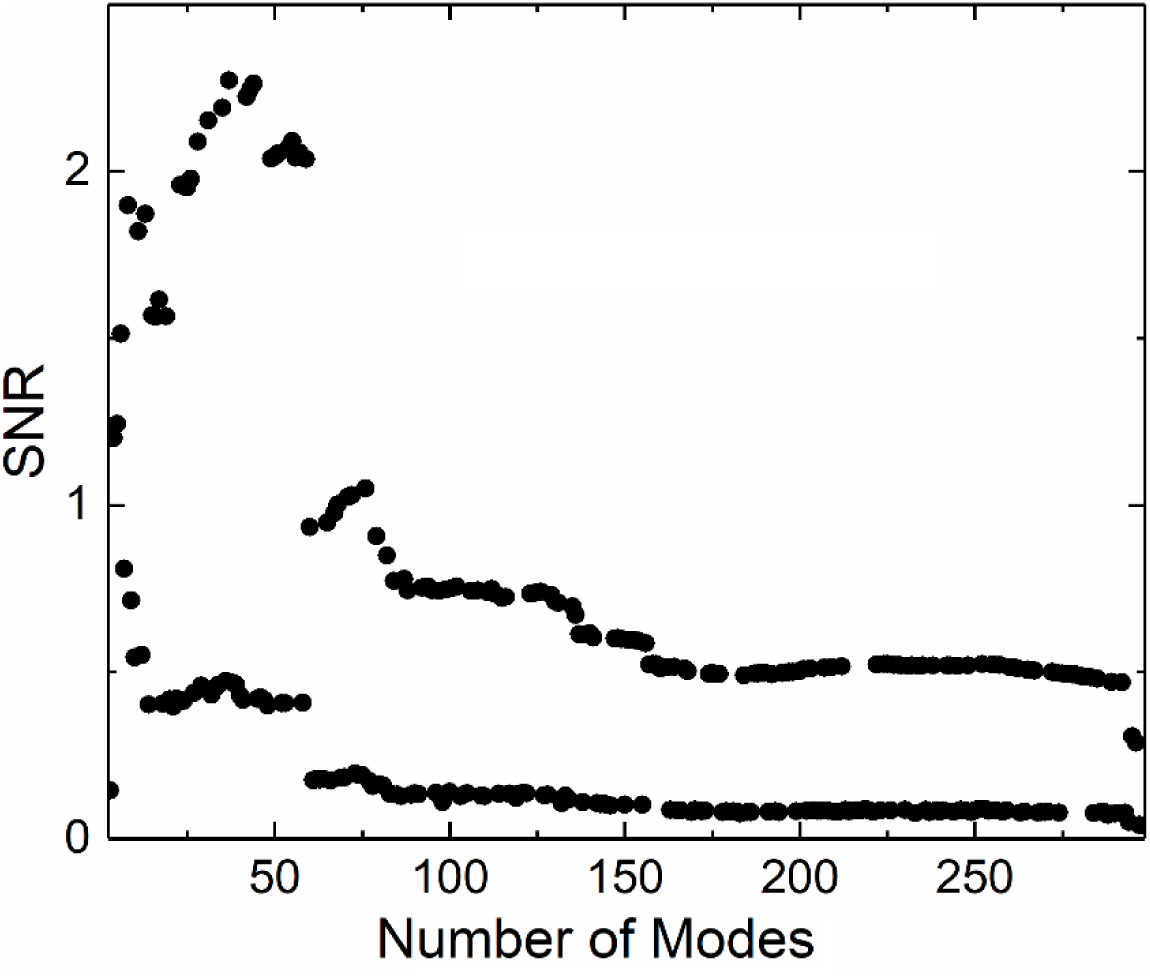
SNR values for PTP1B.

### 3-2. Bovine β-Lactoglobulin

β-lactoglobulin is involved in transmitting allosteric signals especially in the context of biosensors as shown by the experimental work of Yang et al., (Yang, et al., 2024). Changes in the environment around W19, such as ligand binding such as fatty acids or retinol, influence the conformation of E108 and facilitate allosteric communication within the protein. Changes in conformation of E108 either enhances or inhibits the protein’s functionality as a transporter. In this subsection, we search for residues of that affect the interaction between W19 and E108.

In Figure 8 the asterisks denote the conserved residues (Katakura, et al., 1994; North, 1989; Ragona, et al., 1997) which act as complementary or backup residues that are on the path between W19 and E108. R124 and F136 are two residues, synergistic and redundant, respectively, that affect the ligand binding features of β-Lactoglobulin.

**Figure 8.**
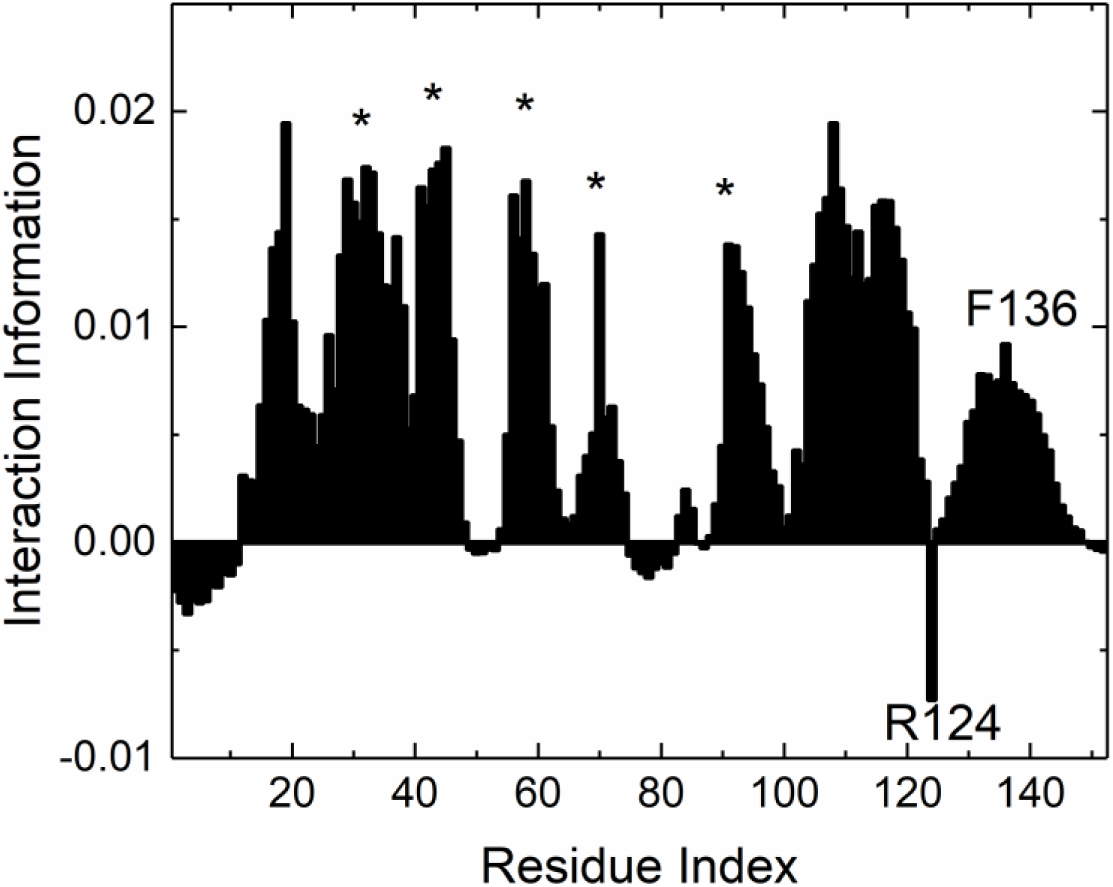
Interaction information profile for Bovine β-Lactoglobulin. The conserved residues denoted by asterisks are backup residues for the W19-E108 interaction.

**Figure 9.**
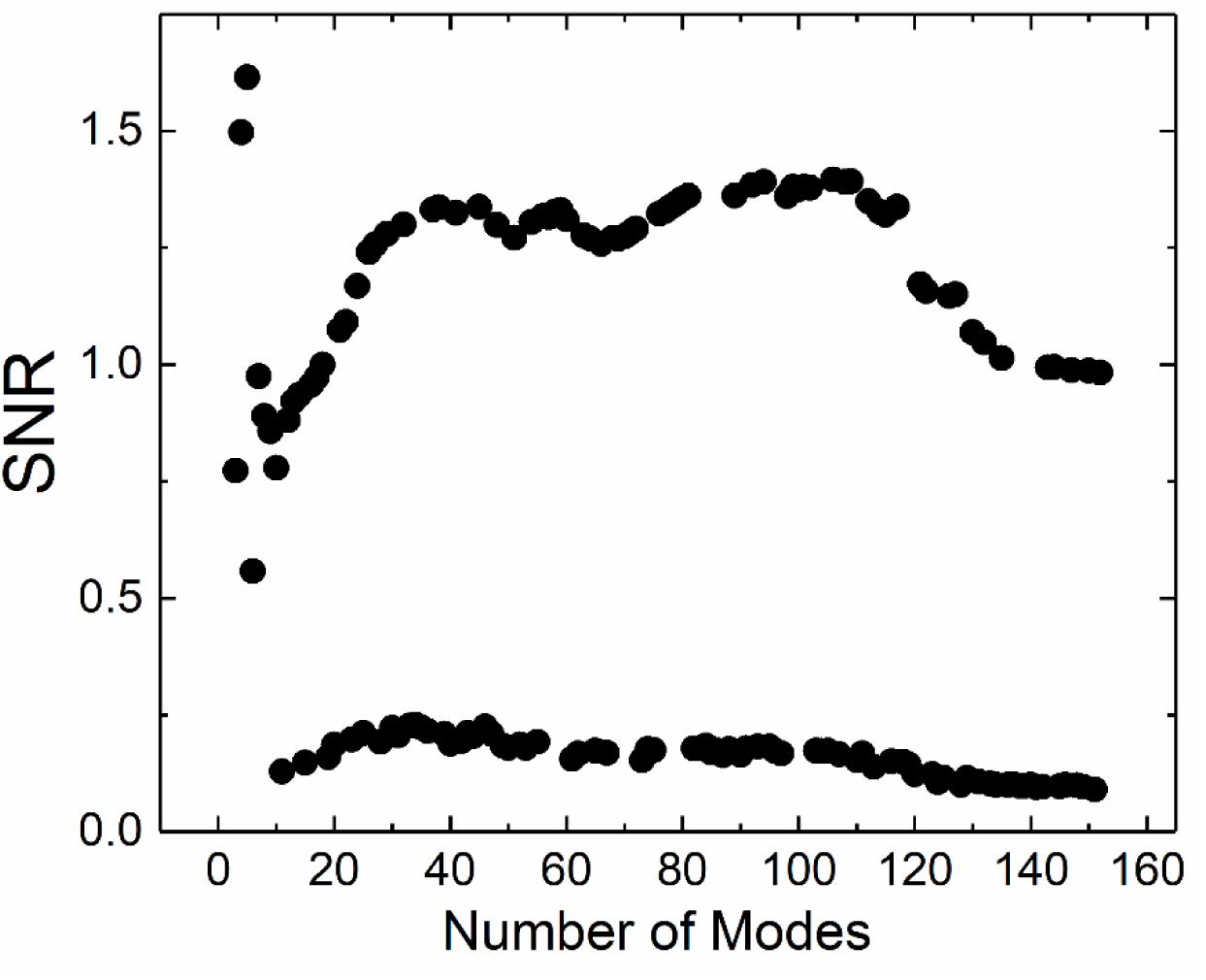
SNR values for Bovine β-Lactoglobulin.

SNR values for β-lactoglobulin are significantly different from those in the two previous cases, with ratios exceeding unity even for large eigenvalues. Additionally, the fluctuations in SNR values with alternating successive eigenvalues are substantial. When the signal remains large irrespective of the number of eigenvalues considered, it indicates a robust system with strong signal characteristics that effectively dominate the allosteric communication.

## 4. Conclusion

This study provides a comprehensive analysis of the role of residue fluctuations in allosteric regulation within proteins, emphasizing the interplay between synergy, redundancy, and noise. By utilizing the Gaussian Network Model and information-theoretic metrics, we have demonstrated how these dynamics contribute to the functional integrity of proteins such as CheY, Tyrosine Phosphatase, and β-lactoglobulin. Our findings show that the first ten modes of fluctuation are particularly critical for maintaining high signal-to-noise ratios, underscoring the necessity of careful eigenvalue selection for accurate predictions of protein behavior.

The analysis indicates that while some eigenvalues correlate with significant displacements, others introduce noise that can obscure meaningful interactions. This highlights the importance of understanding the spectral properties of proteins in relation to their functional dynamics. We have shown that conserved correlations among residues enhance signaling pathways and provide stability against noise-like fluctuations, with redundant residues acting as buffers against potential disruptions caused by mutations or environmental changes.

Additionally, our identification of synergistic interactions among residues aids in mapping allosteric pathways and understanding long-range communication mechanisms within proteins. This work reinforces the notion that proteins should be viewed as adaptive information processing systems rather than static structures, revealing their capacity for efficient and robust communication.

The insights gained from this research have significant implications for future studies on protein engineering and therapeutic interventions aimed at addressing genetic diseases. By advancing our understanding of the complex relationships between residue interactions and their contributions to protein function, we lay the groundwork for innovative strategies in drug design and molecular biology. Ultimately, this study contributes to a deeper understanding of how allosteric regulation operates at a molecular level, paving the way for future research into enhancing protein functionality and stability through targeted modifications.

